# Family permutation profiling identifies a dynamic protein domain as functionally tolerant to increased conformational entropy

**DOI:** 10.1101/840603

**Authors:** Joshua T. Atkinson, Alicia M. Jones, Vikas Nanda, Jonathan J. Silberg

**Affiliations:** Systems, Synthetic, and Physical Biology Graduate Program, Rice University, 6100 Main Street, MS-180, Houston, Texas 77005; Biochemistry and Cell Biology Graduate Program, Rice University, 6100 Main Street, MS-140, Houston, Texas 77005; Center for Advanced Biotechnology and Medicine, Rutgers, The State University of New Jersey, Piscataway, NJ 08854; Department of BioSciences, Rice University, 6100 Main Street, MS-140, Houston, TX, 77005; Department of Bioengineering, Rice University, 6100 Main Street, MS-142, Houston, TX, 77005; Department of Chemical and Biomolecular Engineering, Rice University, 6100 Main Street, MS-362, Houston, TX, 77005

**Author notes:** To whom correspondence should be addressed: Jonathan J. Silberg, Biosciences Department, 6100 Main Street, Houston TX 77005.

**Keywords:** adenylate kinase, circular permutation, combinatorial library, deep mutational scanning, energetic frustration, transposon mutagenesis

## Abstract

To investigate whether adenylate kinase (AK) homologs differ in their functional tolerance to mutational lesions that alter dynamics, we subjected three homologs having a range of thermostabilities to random circular permutation and evaluated where new protein termini were non-disruptive to activity using a cellular selection and deep mutational scanning. Analysis of the positional tolerance to new termini, which increase local conformational entropy by breaking peptide bonds, showed that bonds were either functionally sensitive to cleavage across all three homologs, differentially sensitive, or uniformly tolerant. The mobile AMP binding domain, which displays the highest calculated contact energies (frustration), presented the greatest tolerance to new termini across all AKs. In contrast, retention of function in the lid and core domains was more dependent upon AK melting temperature. Thus, regions of high energetic frustration tolerated increases in conformational entropy in a manner that was less dependent on thermostability than regions of lower frustration. Our results suggest that family permutation profiling identifies primary structure that has been selected by evolution for high frustration that is critical to enzymatic activity. They also illustrate how deep mutational scanning can be applied to protein homologs in parallel to learn how topology and function govern mutational tolerance.

## INTRODUCTION

Protein folding occurs through a funneled energy landscape where the top of the funnel represents the ensemble of unfolded polypeptide conformations, and the bottom represents the folded native-state ensemble with the lowest free energy (1, 2). To encode the native-state ensemble, the amino acids at each native position have been selected through evolution to minimize the average free energy of their contacts. However, some residue-residue contacts can present calculated free energies that are high relative to all other possible residue-residue contacts that could exist at the same location (3). In cases where a given contact has a high energy relative to other possible residues at that location, it has been described as ‘highly frustrated’ (4, 5). Regions of high frustration have been implicated in facilitating protein folding, motion, and binding to other molecules (6–10). These studies suggest that nature selects for high frustration in regions where high conformational flexibility is critical to protein function, and low frustration in regions that are more critical to supporting the stability of the native-state ensemble. While there is evidence that some regions of primary structure are under selection for specific levels of energetic frustration (11, 12), it remains unclear whether selection for residue-residue contacts with low versus high energetic frustration affects tolerance to any specific classes of mutational lesions.

The interplay between stability and dynamics has been intensively studied in the adenylate kinase (AK) family (11–16). During the AK catalytic cycle, a reaction that involves reversible phosphoryl transfer (ATP + AMP ↔ 2 ADP), the lid and AMP binding domains undergo coordinated conformational changes that involve local unfolding (12, 13, 17–19), while the core domain remains more rigid and is thought to determine overall thermostability (16, 20). Rational design studies have shown that modulation of AK dynamics affect the temperatures where maximal activities are observed (13, 16, 19). Mutational studies have also provided evidence that the lid domain may be less tolerant to changes in conformational entropy because the dynamics of this domain have been fine-tuned for substrate binding (19). The AMP binding domain, in contrast, appears more tolerant to enhanced conformational dynamics as local unfolding is thought to control catalytic turnover (19). While these studies have provided insight into the effects of a handful of mutations on AK structure and function in a small number of model systems, it remains unclear how these trends relate to other AK family members with distinct primary structures and thermostabilities.

One challenge with studying the effects of mutations on structure, function, and dynamics across a protein family is the limited throughput of *in vitro* measurements. In contrast, cellular selections represent a simple approach for rapidly assessing the functional tolerance of many mutations in parallel (21–23), although this approach yields more limited biophysical insight. With cellular assays, profiles of functional tolerance can be rapidly generated by creating a library of mutants, using a high-throughput assay to enrich for mutants that are active, sequencing the library of mutants before and after functional analysis, and calculating the relative abundance of each sequence following the functional enrichment (21, 22, 24). Changes in the abundances of each sequence can be used to estimate biological activity of each mutant by calculating a fitness score (25–27). To date, this approach has been applied to a wide range of proteins (21–27). However, the effects of primary structure and protein stability within a protein family has not yet been examined, and it is unclear whether there are conserved patterns of mutational tolerance across protein orthologs that differ in primary structure and stability.

Recently, a method for creating combinatorial libraries was described that alters conformational entropy at different locations in a protein. This approach, which is called circular permutation profiling with DNA sequencing (CPP-seq), randomly samples all sequence permutations in a protein by covalently connecting the original termini and creating new termini elsewhere in the primary structure (28). When proteins are created with this type of topological mutation, the local chain entropy is increased at the location where the new termini are created by cleaving the peptide backbone (29–32). Application of CPP-seq to an AK with extreme thermostability, T_m_ > 99°C (33), revealed that more than half of all possible circularly permuted variants retain biological activity at 42°C (28). While this study identified diverse regions in all three AK domains that are functionally tolerant to changes in conformational entropy that can arise from new protein termini, it remains unclear whether the observed pattern of tolerance depends upon AK sequence, energetic frustration, or thermostability. Because proteins with enhanced thermostability are buffered from other classes of mutations (34–37), it seems likely that AK with lower thermostability will be more sensitive to this class of topological mutation.

To better understand how thermostability influences protein tolerance to circular permutation, we subjected three AK homologs to CPP-seq and used a cellular assay to quantify protein fitness at a fixed temperature. We targeted AKs with a range of thermostabilities and reported crystal structures. These experiments revealed that the fraction of functional variants in each library correlates with parental protein thermostability. At the residue-level, we found that a large fraction of the permutated proteins were either non-functional across all three AKs or displayed fitness that correlates with AK thermostability. However, a subset of circularly permuted AK retained fitness across all three variants, and some presented higher fitness than the parental proteins. A comparison of the positional tolerance to permutation revealed that this latter group of variants arose from the creation of new protein termini at native sites within the AMP binding domain, which has contacts with consistently high energetic frustration across all three AKs. This correlation suggests that energetic frustration may be useful for predicting regions of homologous proteins that are functionally tolerant to increases in conformational entropy arising from circular permutation.

## RESULTS

### Library construction and characterization

A major challenge with using cellular assays to compare the mutational tolerance of protein orthologs is achieving uniform expression in cells, such that the retention of function only depends upon protein stability and total activity. This is particularly challenging with topological mutations like circular permutation where the mutation alters the genetic context of the ribosomal binding site (RBS) that controls translation initiation (38). In a previous study, we showed that transposon mutagenesis can be used to build vector libraries that express different circularly permuted variants of any protein with the same translation initiation rate (39). With this approach, which is called PERMUTE, libraries of permuted genes are created by randomly inserting a linear vector, called a permuteposon (40), into a gene that has been circularized by connecting the first and last codons using a non-disruptive linker sequence. In cases where permuteposon P1 is used to build libraries (41), the resulting vectors express different permuted proteins with the same 18 amino acid peptide fused to the N-terminus (40). The DNA encoding this peptide maintains the genetic context of the RBS so that translation initiation is constant across different protein variants (39).

Three different AKs were subjected to random circular permutation using PERMUTE (28), including *Bacillus globisporus* AK (*Bg*-AK), *Bacillus subtilis* AK (*Bs*-AK), and *Geobacillus stearothermophilus* AK (*Gs*-AK). These AK orthologs were chosen because they exhibit melting temperatures that span over 30°C (42, 43). A sequence alignment reveals that these AKs are identical in length and all contain the Cys-X_2_-Cys-X_16_-Cys-X_2_-Cys/Asp zinc-binding motif (residues 130-153) in their lid domain (Figure S1), which is typical of AKs from gram-positive bacteria (43). Additionally, these AKs exhibit high pairwise sequence identities (66 to 74%), and structures with low RMSD (0.73 to 1.76Å) (42). Because their native termini are proximal, all of the circularly permuted AKs were expressed with their native termini linked using a tripeptide (Ala-Ala-Ala). This peptide was previously found to be compatible with creating functional permuted *Thermotoga neapolitana* AK (*Tn-*AK) (40).

In total, each AK library contains up to 1320 different vectors. This diversity is produced because PERMUTE randomly inserts the permuteposon at every location in the AK genes (651 base pairs) that were circularized using a 9 base pair linker. Half of the insertions (n=660) into the circular genes occur such that the regulatory elements that drive expression in the permuteposon are parallel (P) with the open reading frames (ORFs) encoding each circularly permuted gene (Figure 1A). In contrast, the other half are in an antiparallel (AP) orientation and cannot express circularly permuted genes. Among the P variants, only one third of the permuted genes are in frame (n=220), and only 216 of these express permuted proteins. Four of the variants occur within the linker and express a full length AK, which serve as a frame of reference for parent protein fitness in all experiments.

**Figure 1.**
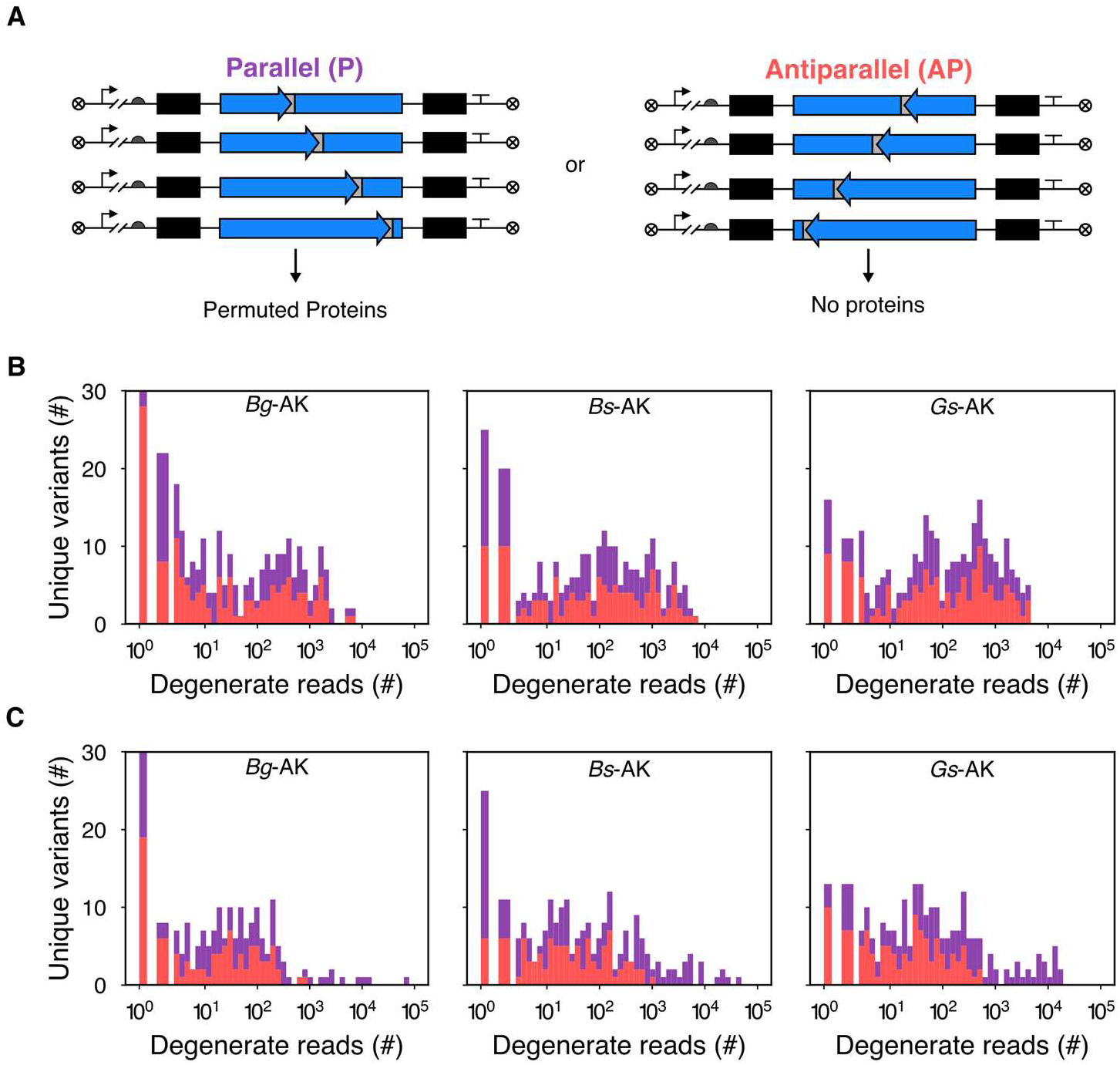
Permuted gene abundances versus gene orientation. (*A*) Generation of libraries using PERMUTE requires mixing a circular gene, a permuteposon, and MuA transposase. The permuteposon inserts in two orientations with equal frequencies (28). The orientation where the regulatory elements *i.e.*, promoter, RBS, and terminator, and the permuted genes are in the same direction is designated as *parallel*. Conversely, when the permuteposon inserts in the opposite direction such that the regulatory elements and the permuted genes are oriented in opposite directions this is designated as *antiparallel*. Only when the permuteposon inserts in the *parallel* orientation does it generate a vector capable of expressing a circularly permuted protein with an 18-residue peptide amended to the new N-terminus. For each (*B*) unselected library and (*C*) selected library we evaluated the number of degenerate, in-frame sequences at each position in both the P (purple) and AP (red) orientations.

We transformed all three libraries into *E. coli* DH10B and quantified the number of unique colony forming units (cfu) following an overnight incubation under non-selective conditions. This analysis revealed that the libraries sampled more than 174000 (*Bg*-AK), 164000 (*Bs*-AK), and 28000 (*Gs*-AK) unique vectors containing AK genes. To evaluate the sequence diversity created, each library was purified, barcoded, and subjected to deep sequencing using MiSeq (28). This analysis yielded more than three million total sequence reads for each library. Table 1 lists the abundances for the P and AP reads in each library. The numbers of desired expression vectors sampled, *i.e*., those that are in frame and parallel, were 52332 (*Bg*-AK), 67193 (*Bs*-AK), and 70258 (*Gs*-AK).

**Table 1.** Parallel and antiparallel sequence read counts in each library. The data generated by MiSeq analysis, including the total reads having the barcode corresponding to each AK ortholog in both the unselected and selected libraries, the number of reads with a permuteposon barcode and 11 base pairs (bp) of AK gene used to identify the location of the new termini. For reads with permuteposon barcode and 11bp of AK gene, we list the number of P and AP that are in frame, +1 frame, and −1 frame. The ratio of selected to unselected abundances (S/US ratio) is also shown.

|  | *Bg*-AK |  |  | *Bs*-AK |  |  | *Gs*-AK |  |  |
| --- | --- | --- | --- | --- | --- | --- | --- | --- | --- |
|  | Unselected | Selected | S/US Ratio | Unselected | Selected | S/US Ratio | Unselected | Selected | S/US Ratio |
| Total reads | 3345761 | 2943111 | 0.88 | 3541540 | 3583510 | 1.01 | 3486607 | 2872462 | 0.82 |
| +11bp AK | 224235 | 245313 | 1.09 | 230619 | 248489 | 1.08 | 232916 | 224023 | 0.96 |
| P in frame | 52332 | 224028 | 4.28 | 67193 | 226735 | 3.37 | 70258 | 208547 | 2.97 |
| AP in frame | 48893 | 5699 | 0.12 | 67747 | 9361 | 0.14 | 67197 | 7003 | 0.10 |
| P +1 frame | 49708 | 6422 | 0.13 | 34633 | 4179 | 0.12 | 32411 | 2544 | 0.08 |
| AP +1 frame | 48340 | 5380 | 0.11 | 34362 | 4675 | 0.14 | 33684 | 3259 | 0.10 |
| P -1 frame | 12583 | 2344 | 0.19 | 13191 | 1820 | 0.14 | 14634 | 1264 | 0.09 |
| AP -1 frame | 12379 | 1440 | 0.12 | 13493 | 1719 | 0.13 | 14732 | 1406 | 0.10 |

When creating libraries using transposon mutagenesis, the abundances of each permuted gene can vary widely (28), although the relative abundance of P and AP variants of each permuted gene is typically similar. Figure 1B shows that relative abundance of each permuted gene varied by up to three orders of magnitude within each library. In contrast, the abundances of P and AP variants arising from insertion at the same location presented abundances that differed by <7% (Figure S2). The variants with the highest abundances in each library were different. With the *Bg*-AK library, the permuted gene that started with the Pro-165 codon was most abundant (n=7307), while the AK genes that started with codons encoding Val-115 (n=5039) and Thr-31 (n=4456) were most prevalent in the *Bs*-AK and *Gs*-AK libraries, respectively.

### Thermostability correlates with mutational tolerance

Our goal was to identify permuted AKs in each library that retain catalytic activity at 42°C. To achieve this goal, we used a high-throughput cellular assay to detect biologically-functional AKs. To profile which variants retain AK function in cells, each library was enriched for vectors that express an active AK using bacterial complementation with *E. coli* CV2 (44), a strain with a temperature-sensitive AK that cannot grow above 40°C unless a functional AK is expressed in trans (45). The vector ensembles were purified following the selection and deep sequenced, and retention of function was quantified by analyzing the ratio of P and AP sequence abundances before and after selection as previously described (28). By quantifying the ratio of P to AP variants of each permuted AK, we were able to identify variants that were enriched to a similar extent as the parental AKs, *i.e.*, those with similar total activity, as well as variants that are enriched to a greater extent than the inactive AP variants.

Following selection of each library for functional AKs using *E. coli* CV2, we quantified the number of unique vectors sampled. All three libraries yielded >48000 unique vectors. Table 1 shows that MiSeq analysis of the selected reads yielded similar total counts as the unselected libraries. Figure 1C shows that genes with the highest abundances in the selected libraries were in the P orientation. Across the selected libraries, the fraction of in frame P vectors (0.968±0.006) was 30-fold greater than the AP fraction (0.032±0.006). This result can be contrasted with the unselected libraries, where the in frame P and AP fractions were similar (0.509±0.008 and 0.491±0.008, respectively). These results illustrate how the selective pressure applied by the cellular assay dilutes the AP variants that cannot express a permuted AK relative to the P variants that can express active permuted AKs.

To establish which permuted AKs are active in *E. coli* CV2, we calculated the fold change in abundances of each unique permuted AK variant in the selected and unselected libraries in both the P and AP orientations. In PERMUTE libraries, cognate P and AP variants are synthesized at similar proportions (Figure S2), and thus, represent paired data whose ratios can be used to assess the significance of sequence enrichment across the wide range of initial abundances observed (28). While the P variants express the different permuted proteins, the AP variants are unable to express proteins and cannot be enriched by the selection. In this way, the AP variants serve as an internal frame of reference for evaluating whether their cognate P variants are biologically active (enriched more than AP variants) or inactive (similarly diluted as AP variants).

We used a negative binomial distribution to model the mean and the variance of the fold change of AP variants in the selected and unselected conditions, *i.e.,* 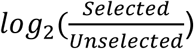 relative to each variant’s initial abundance in the unselected library (28). Using these values, we investigated which P variants are biologically active by identifying those that were significantly enriched by the selection (two-tailed t-test, *P* < 0.01). With this analysis (Figure S3), 25.8% (n=38/147) of the sampled P variants were significantly enriched in the *Bg*-AK library, while 43.9% (n=65/148) and 59.7% (n=86/144) were significantly enriched in *Bs*-AK and *Gs*-AK libraries, respectively. Comparing the fraction of functional variants with T_m_ reveals a linear relationship (Figure 2A, r^2^=0.82). This trend shows that thermostability buffers AKs from the destabilizing effects of circular permutation, similar to the buffering effects observed with other classes of mutations (34–37). This buffering is thought to occur because circular permutation has similar destabilizing effects on structurally-related AK homologs (Figure 2B), yielding a similar ensemble of ΔΔG_f_ in each topological mutant library.

**Figure 2.**
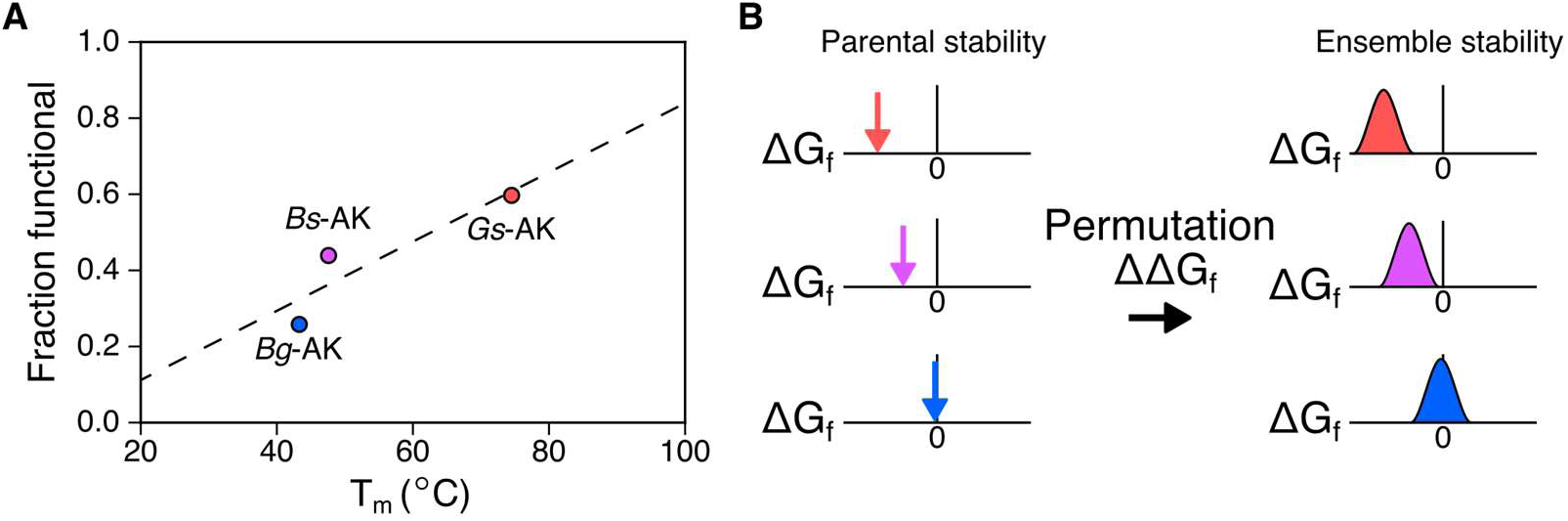
Thermodynamic stability correlates with tolerance to permutation. (*A*) The fraction of P variants sampled in the libraries that were biologically active following selection is plotted relative to the melting temperature of each AK parent. A linear fit (y = 0.01x - 0.07, R^2^ = 0.822) is shown as a dashed line. (*B)* This trend suggests that circular permutation has similar destabilizing effects on structurally-related AK homologs yielding a similar ensemble of ΔΔG_f_.

### Family-level fitness of topological mutants

To determine if functional tolerance to new termini is dependent upon domain location, we next compared the enrichment of each in frame P variant with AK structure (Figure 3). This analysis reveals that new termini are differentially tolerated at diverse locations within the different AK orthologs. Circularly permuted AK were uniformly inactive when new termini were generated within the glycine rich p-loop (residues 7-15), the region of the core domain that constitutes the active site in AKs and other related kinases (42, 46). In contrast, the lid (residues 128-159) and core (residues 1–30, 60–127, and 160–217) domains both varied in their tolerance to new termini across the different AK orthologs, with some positions being uniformly intolerant and other positions exhibiting tolerance that varied across the orthologs. Surprisingly, a large fraction of the permuted AK having new termini within the mobile AMP binding domain (residues 31-59) presented biological function in all three AK homologs. In *Escherichia coli* AK *(Ec*-AK), this domain uses local unfolding to control the rate limiting step of the catalytic cycle, product release (12, 19).

**Figure 3.**
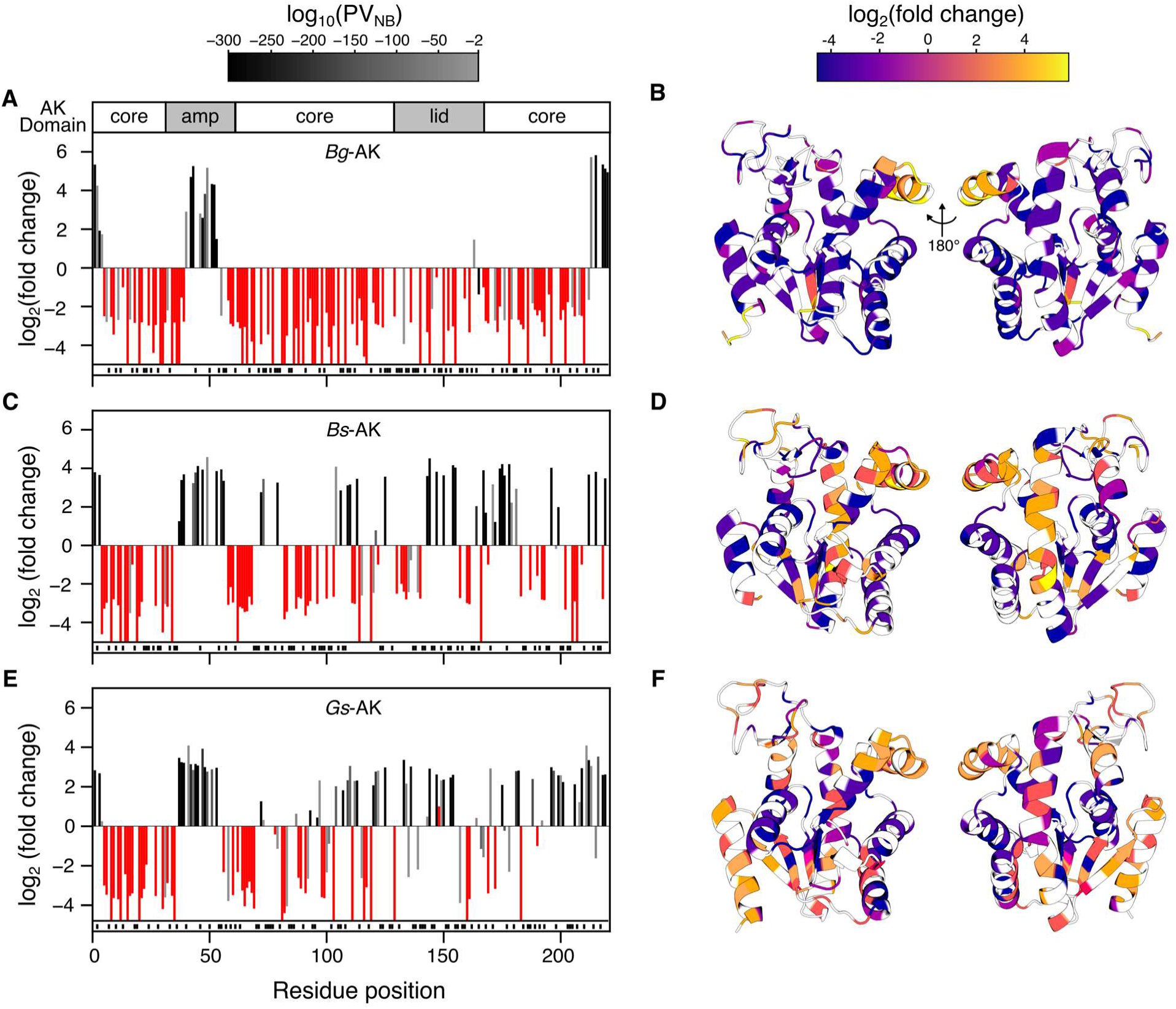
Comparison of circular permutation profiles of each AK. For each P variant, the log_2_(fold change) is shown as a function of the AK residue found at the N-terminus of the circularly permuted protein derived from (*A*) *Bg*-AK, (*C*) *Bs*-AK, and (*E*) *Gs*-AK. *P*-values obtained from the negative binomial model (PV_NB_) are color coded with values >10^−2^ in red and variants having values ≤10^−300^ in black and those variants displaying intermediate values shaded as indicated by the bar. The AK domain structure is shown at the top as a frame of reference. Variants no longer observed following selection are shown as bars that reach the line at the bottom of the graph. The cognate P and AP variant pairs absent from both the unselected and selected libraries (n = 73, 72, and 76 for Bg-AK, Bs-AK, and Gs-AK, respectively) are indicated as black lines shown below the x-axis. The log_2_(fold change) is compared with AK structure for (*B*) *Bg*-AK (PDB: 1S3G), (*D*) *Bs*-AK (PDB: 1P3J), and (*F*) *Gs*-AK (PDB: 1ZIO) with the residues colored according to their log_2_(fold change) as indicated by the bar. Unsampled positions are in white.

In our libraries, there are two frames of reference for biological activity: (1) AP variants that cannot express AKs, which are diluted by the selection (Figure S3), and (2) native AK, whose enrichment is dependent upon total activity. Because each AK library was selected in a different experiment, we evaluated the relative enrichment of each parental AK, which was encoded and observed in all three libraries following the selections. This analysis revealed that the fold change of each native AK decreases linearly (r^2^ = 0.99) as the fraction of functional variants in each library increases (Figure S4). This finding illustrates the additional selective pressure of increased competition that arises as the fraction of functional variants competing with native AKs changes in each experiment. To enable us to compare the sequence enrichment trends across the three libraries, we converted the fold enrichment value into a measure of parental fitness by normalizing the enrichment value for each permuted variant to that observed with the native AKs. Due to partial sampling in the individual libraries, only 38.7% (n = 84/217) of the circularly permuted AKs were observed in all three libraries and thus available for family permutation profiling. Those circularly permuted AKs sampled in all three libraries were distributed across the AMP binding, lid, and core domains.

To first evaluate how the fitness of each trio of the permuted AK homologs relates to the location of their protein termini, we compared the fitness of each variant to the number of AK orthologs in which that position was observed to be significantly enriched. With this family-level analysis, we observed different trends when looking at homologous circular permuted variants derived from the different parental AKs (Figure 4A). Some permuted AK presented low fitness values across all three AK homologs that could not be distinguished from cells lacking an AK. This trend indicates that these positions are uniformly intolerant to permutation across all three AK homologs. Other permuted AK presented fitness values that varied across the different parents. At some locations, we observed one out of the three structurally-related variants as enriched while two out of three of the variants were enriched at other locations. Surprisingly, some AK variants presented parent-like fitness values across all three AK homologs.

**Figure 4.**
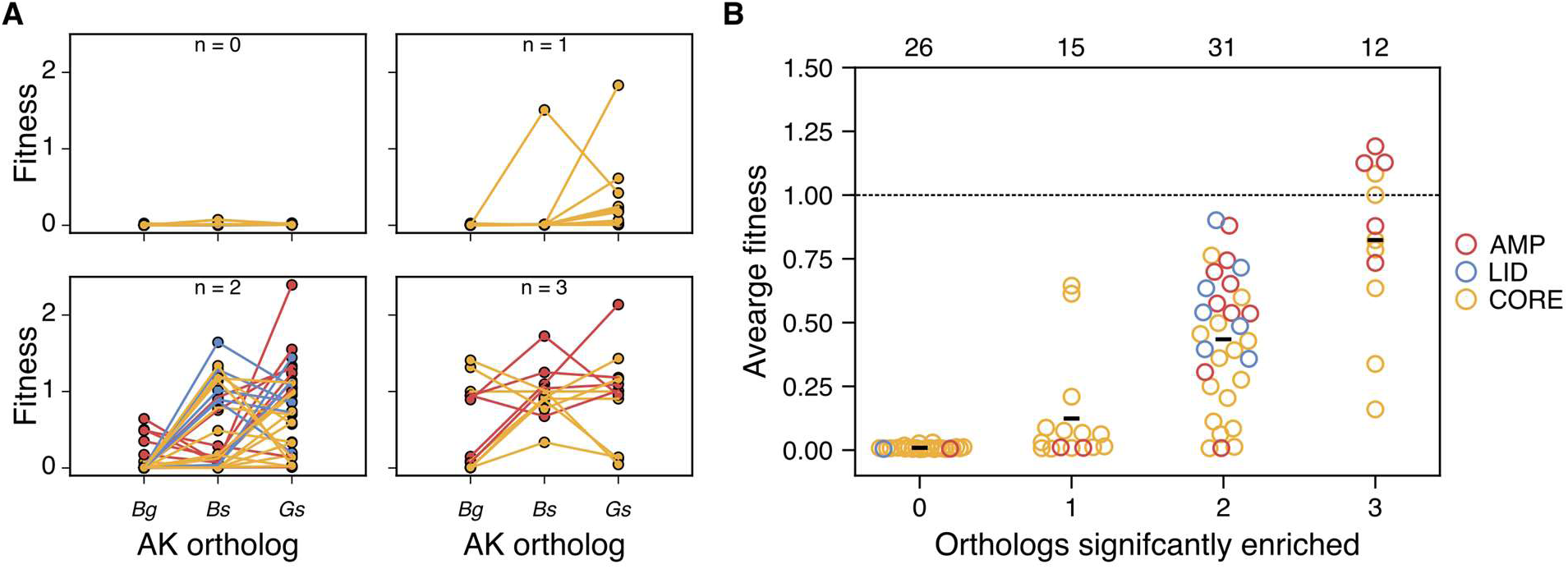
Comparison of family enrichment and fitness. (*A*) For variants that were sampled in all three AK libraries (n = 84), the fitness score (fold change relative to native AK) is shown. In each box, the number of AK homologs across the three libraries that were found to be significantly enriched using PV_NB_ is noted as 0, 1, 2, or 3. Data for each topological variant is connected by a line to illustrate the trend across homologous permuted proteins derived from parent AKs having distinct thermostabilities. Those topological mutants that were uniformly biologically active across all three libraries (n = 3) presented a wide range of fitness values (ranging from 0.004 to 2.13). (*B*) For the same set of circularly permuted protein homologs, the average fitness of all three orthologs is compared to the number of orthologs found to be significantly enriched using PV_NB_. The number of data points in each bin is noted at the top. The average fitness of all the orthologs in a column is noted with the short black line, while the average fitness of the native AKs are indicated by the dashed line. The domain location of the new protein termini in each permuted protein trio is indicated by the color of the ring according to the legend.

To visualize all of this family-level data, we analyzed the *average* fitness of each circularly permuted variant observed in the *Bg-*AK, *Bs-*AK, and *Gs-*AK libraries (Figure 4B). This analysis revealed that circularly-permuted proteins that were inactive in all three libraries primarily arose from creation of new protein termini mostly in the core domain, although variants having protein termini in all three domains were observed. In contrast, those circularly permuted variants that were active in all three libraries arose from new termini in either the AMP binding or core domains. The variants arising from protein termini in the AMP binding domain exhibited the highest average fitness among these variants. Permuted AKs that were significantly enriched in either one or two orthologs contained protein termini in all three domains.

### Relationship between fitness and energetic frustration

Because circular permutation creates increased conformational flexibility at the location of the new termini, we hypothesized that new termini might be tolerated to a greater extent at locations that have been selected to support local flexibility. One way that such locations have been identified is by identifying the residue-residue contacts in a protein that exhibit higher pairwise energies (frustration) than other amino acids at that same position. To evaluate how frustration changes across each domain within the two conformations, we used the frustratometer server to generate profiles of inhibitor-bound and substrate-free *Ec*-AK (Figure 5A) (47). At native locations where the amino acids make residue-residue contacts that are lower energy compared to all other amino acid possibilities at that contact, the profile yields a low frustration value. In contrast, native locations having amino acids that present higher contact energies compared with all other theoretical amino acid combinations at that particular contact yield high frustration values. In the inhibitor-bound closed conformational state, there is a shift in this energetic profile with an increase in the density of highly-frustrated contacts within the AMP binding domain and a subtle decrease in the lid domain. These state-specific frustrated contacts are thought to facilitate the local unfolding events in the AMP binding domain that are coupled to the lid opening at the end of the AK catalytic cycle (11). To determine if the AKs used to build our libraries also exhibit high frustration in the AMP binding domain, we calculated profiles of their energetic frustration using inhibitor-bound AK structures (Figure 5B). These calculations revealed that all three proteins exhibit similar patterns of energetic frustration as the closed conformation of *Ec*-AK, even though their thermostability varies widely. This result suggests that this pattern of energetic frustration has been selected during evolution to support catalysis across different ranges of temperatures. Comparing these profiles with AK structure reveals that regions of high energetic frustration are localized in the AMP binding and lid domains as well as surface exposed sites in the core domain (Figure 5C). In contrast, regions of minimal energetic frustration are localized primarily within the core domain. These residue-residue contacts are thought to play an important role in controlling protein folding and maintaining a preorganized active site that is poised for catalysis (15). Together these suggest that AKs have selected residue-residue contacts with specific levels of frustration at individual native sites to support protein folding, substrate binding, and conformational dynamics critical to catalysis (Figure 5D).

**Figure 5.**
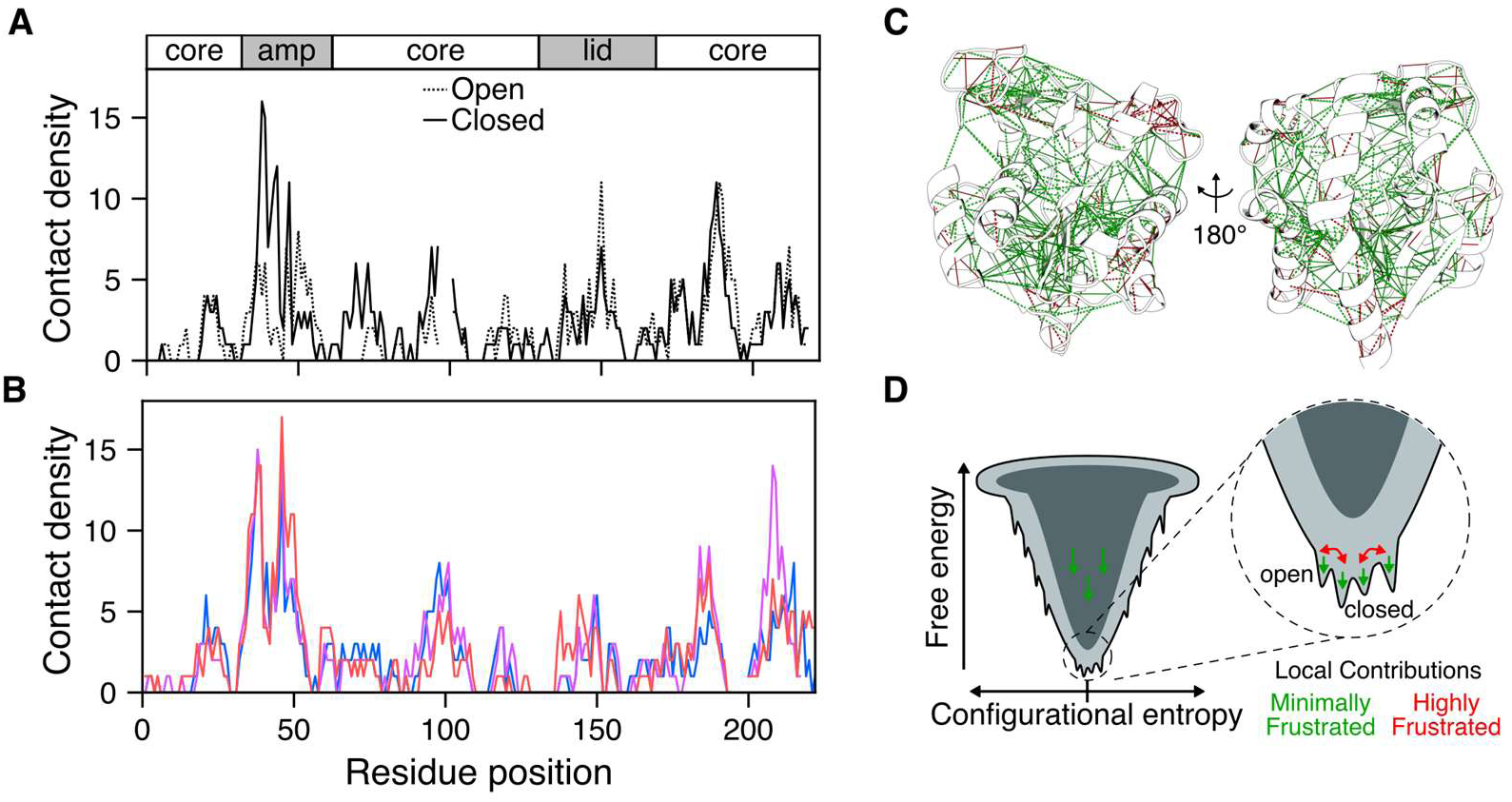
Energetic frustration of intramolecular contacts in adenylate kinase. (*A*) Frustratograms showing the calculated configurational frustration of the open (dashed) and closed (solid) conformations of *Ec-*AK (calculated from PDB: 4AKE and 1AKE, respectively). The density of contacts in a 5Å sphere with highly frustrated contact energies is plotted at every AK residue position. (*B*) A comparison of the frustratograms of the closed conformations of *Bg*-AK (blue), *Bs*-AK (purple), and *Gs*-AK (red). (*C*) The contact energy frustration is mapped onto the tertiary structure of a closed structure (PDB: 1AKE). Highly frustrated contacts are displayed in red, while minimally frustrated are shown in green. Direct contacts are solid lines, while water mediated contacts are dashed lines. (*D*) Influence of contact frustration on the global protein folding and functional energy landscape. Minimally frustrated contacts promote protein folding into the low-energy native ensemble, while within this ensemble highly frustrated contacts facilitate the sampling of conformations including those important for catalysis.

We next investigated how the locations of the termini in variants having different levels of fitness relates to primary structure, tertiary structure, and energetic frustration (Figure 6A-B). This comparison revealed that the region of the AMP binding domain where new termini are functionally tolerated to the greatest extent overlaps with the region containing the highest density of frustrated contact energies. When looking at the positions within the AMP binding domain that are enriched across either zero, one, two, or three AK orthologs (Figure S5), the average fitness increases as the energy landscape becomes more frustrated. Like the AMP binding domain, regions proximal to the native protein termini exhibit functional tolerance to new termini across all three AKs, even though this region does not exhibit high frustration like the AMP binding domain. Similarly, a patch within the middle of the α7 helix that is surface exposed on the back of the core domain is uniformly tolerant to new termini, even though this region exhibits moderate frustration across all three AK homologs.

**Figure 6.**
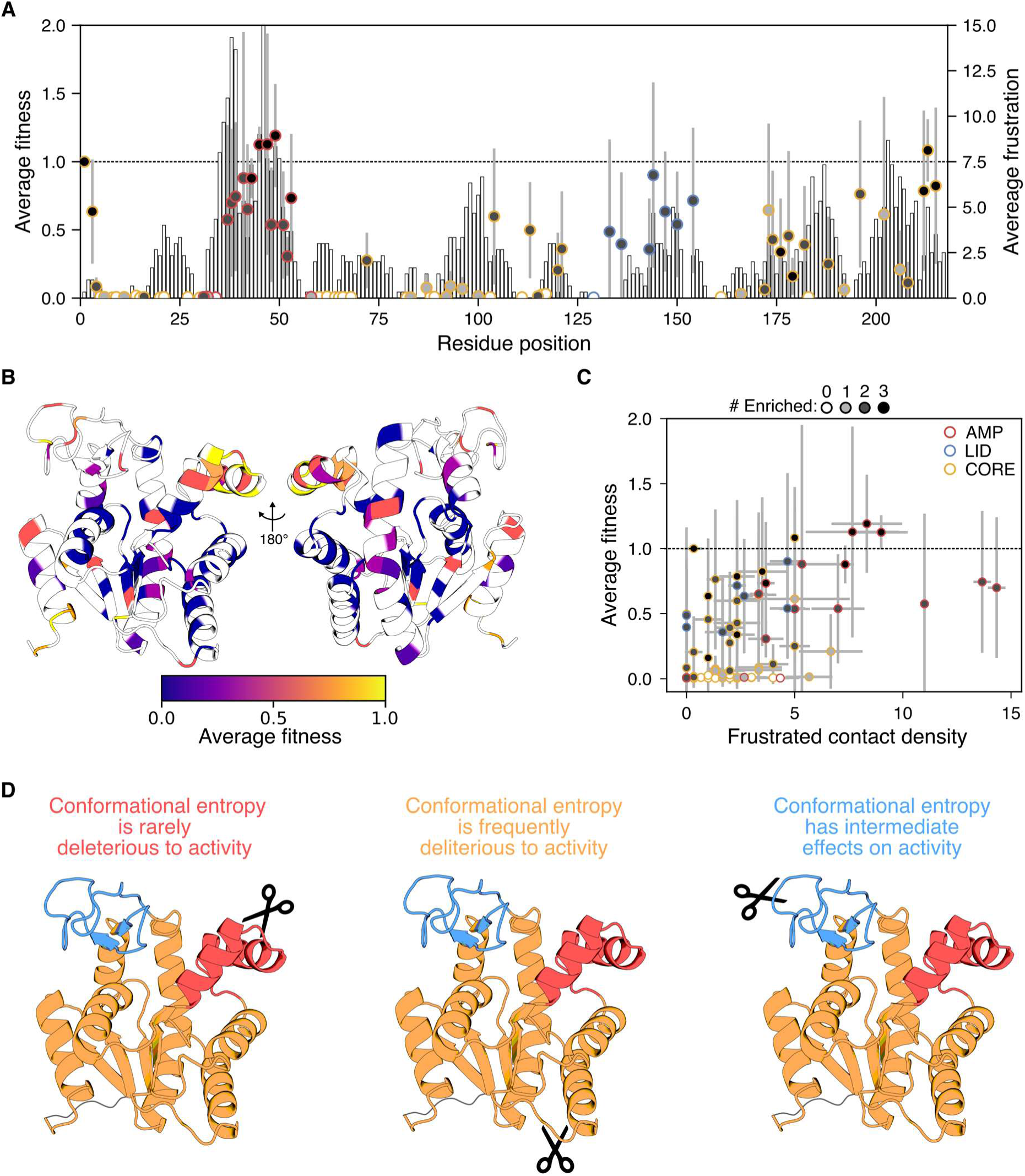
Comparison of family permutation profiles and energetic frustration. (*A*) For topological mutants that were sampled in all three libraries (n = 84), the average fitness and average frustration are compared to the position in the primary structure. The circles represent the average fitness of each variant with the standard deviation shown in gray. The shading of each circular data symbol represents the number of orthologs in which the variant was found to be enriched. The bars represent the average density of highly frustrated contacts at each position observed across all three structures. The dotted line represents the native AK fitness. (*B*) The average fitness of all three orthologs is compared to the tertiary structure (PDB: 1S3G). The color of the residue corresponds to the average fitness of a permutation at each site according to the color bar. Positions that were unobserved in one or more orthologs are shaded white. (*C*) The average fitness of each trio of homologous permuted AK is compared to the average density of highly frustrated contacts at that position in the native AK structures. Each circle represents the average fitness and the average frustration observed across all three structures. The standard deviations are shown in gray. The shading of each circular data symbol represents the number of orthologs in which the variant was found to be enriched. The dotted line in all four panels represents the native AK fitness observed in each library. Spearman’s rank correlation coefficient between the fitness and frustrated contact density was 0.512 with a two-tailed *P-*value = 6.13 × 10^−7^. (*D*) Higher conformational entropy caused by permutation in the AMP binding domain is not deleterious to enzyme fitness, while permutation in the core and lid domains negatively impacts enzyme fitness.

Because the AMP binding domain exhibits high tolerance to permutation across all three AKs, we next compared the average fitness of each variant with the average density of highly frustrated contacts found in the ‘closed’ state structures of each AK ortholog (Figure 6C). This analysis revealed a significant correlation (Spearman’s rank correlation *R_SR_* = 0.512, two-tailed *P-*value = 6.13 × 10^−7^) between the density of highly frustrated contacts and fitness across the different permuted variants. The circularly permuted AKs that were not significantly enriched (fitness = 0.009 ±0.006) across all three libraries display the lowest average densities of frustrated contact densities (1.44 ±1.33 contacts). These topological mutants primarily arise from permutation that creates new termini in the core domain. In contrast, those variants that were uniformly enriched across all three AK libraries (fitness = 0.823 ±0.309) present the highest average densities of frustrated contacts (4.29 ±2.97 contacts). These topological mutants have new termini that are primarily localized in the AMP binding domain. The variants only enriched in one or two AK orthologs display intermediate values of average fitness (0.124 ±0.205 and 0.436 ±0.254, respectively) and frustrated contact densities (2.55 ±1.97 and 3.35 ±3.68, respectively). This latter group of topological mutants arise from new termini in all three domains.

To further explore how protein dynamics influence tolerance to permutation we compared average fitness to several other experimental and computational metrics. Comparison of the average fitness with normalized B-factors from the inhibitor-bound crystal structures of the three orthologs (42) revealed a significant correlation (Figure S6A, Spearman’s rank correlation *R_SR_* = 0.439, two-tailed *P-*value = 2.95 × 10^−5^), although the P-vlaue for this correlation was lower than that observed with frustration. Comparison of the average fitness with nuclear Overhauser effect (NOE) data from nuclear magnetic resonance (NMR) studies of *Ec*-AK (48) revealed an inverse correlation (Figure S6B, Spearman’s rank correlation *R_SR_* = −0.324, two-tailed *P-*value = 6.00 × 10^−3^). To take an entropic perspective, we compared the average fitness with predictions of residue-level conformational entropy made by POPCOEN, a neural-network trained on molecular dynamics simulations of ~1000 proteins (49). This analysis also revealed a significant correlation (Figure S6C, Spearman’s rank correlation *R_SR_* = 0.380, two-tailed *P-*value = 3.71 × 10^−4^). However, when we compared average fitness to the residue level predictions of conformational entropy loss upon folding from the denatured state ensemble to the native state ensemble using the calculator PLOPS (50) no significant correlation was observed (Figure S6D).

To establish how the different protein dynamics metrics relate to one another, we evaluated the pairwise relationship of each metric. These comparisons revealed significant positive correlations between B-factors and the density of highly frustrated contacts calculated using the frustratometer as well as B-factors and the residue-level conformational entropy calculated by POPCOEN (Figure S7A); B-factors did not correlate with the entropy loss calculated by PLOPS. In addition, inverse correlations were revealed between NOEs and both frustration and POPCOEN calculated entropies (Figure S7B). However, no significant correlation was observed between NOEs and PLOPS. Furthermore, no significant correlations were observed when comparing the three computational metrics or NOE and B-factors.

## DISCUSSION

Deep mutational scanning is increasingly used to analyze the contributions that individual residues make to protein function (22, 24, 51). This approach has identified residues that underlie protein solubility (52), protein-protein interactions (53, 54), membrane protein insertion (55), thermostability (56–58), substrate specificity (59), enzymatic function (56), and mutational epistasis (60). Such efforts have provided fundamental insight into sequence-function relationships within individual proteins. By analyzing the effects of mutations on protein functions across different conditions, deep mutational scanning has shown that environmental conditions can affect how mutations contribute to fitness (56, 59, 61, 62). However, the extent to which the results from any individual study are generalizable to structurally- and functionally-related protein homologs is not known. The family permutation profile generated through our study identified sequence-function relationships that are consistent across multiple protein homologs. These results suggest that family mutational profiles can be used to glean permutation rules for protein homologs that exhibit similar biological activities but differ in sequence and stability.

In our experiments, we chose to vary AK thermostability, while assessing protein function at a single temperature. Previous studies have shown that thermostability alters rates of evolution (63) and buffers proteins from the disruptive effects of random amino acid substitutions (34, 37) and backbone fission (64), suggesting that it should be generalizable to topological mutations like circular permutation that alter local conformational entropy. Our measurements extend this trend to circular permutation by showing that the fraction of functional AK increases linearly with the T_m_ of each AK homolog, from 25.8% (*Bg*-AK) to 43.9% (*Bs*-AK) to 59.7% (*Gs*-AK). This observation is consistent with a previous study examining tolerance to permutation in *Tn*-AK, an AK with even higher thermostability (T_m_ = 99.5°C) than the homologs studied herein. Using the same cellular assay, this prior study found that new termini were tolerated in 65.5% of the sampled positions within circular permuted *Tn*-AK (28).

Using AK homologs of varying thermostability for mutational studies at a single temperature is akin to altering the temperature of a reaction being studied with a single enzyme (65, 66). Those proteins that evolved to function closest to the assay temperature typically exhibit enhanced flexibility compared to the more thermostable variants (35, 36). This additional mobility is thought to allow these proteins to sample more conformations in the native basin of the energy landscape resulting in lower occupancy of the functionally folded state, while more rigid thermostable variants occupy a narrower region of the energy landscape increasing the fraction of functionally folded enzyme (35, 36, 67–70). Mutations that locally perturb this flexibility/rigidity tradeoff can have different fitness effects depending on their domain locations (71, 72). This tradeoff has been observed in chimeras created by recombining *Bs*-AK (T_m_ = 47.6°C) and *Gs*-AK (T_m_ = 74.5°C) (16). A chimera made up of the *Gs*-AK core domain and *Bs*-AK AMP binding and lid domains exhibited stability like *Gs*-AK but activity that is higher than both parental proteins (16). In contrast, a chimera having the *Bs*-AK core domain and *Gs*-AK mobile domains displays stability like *Bs*-AK and activity that is lower than both parent proteins (16).

Our results suggest that circular permutation profiling across enzymes exhibiting a range of stabilities represents a simple way to systematically perturb flexibility and identify regions that have been selected to promote folding stabilization versus conformational flexibility. Permuted AKs having protein termini within the core domain presented fitness that is largely dependent upon thermostability. In contrast, permuted AK having protein termini within the mobile AMP binding domain frequently exhibited parent-like fitness across all three AK homologs, and thus, was independent of protein thermostability. The lid domain presented tolerance to protein termini that was intermediate to that observed with the core and AMP binding domains. These findings suggest that there is a benefit to using thermostability as a variable when performing family permutation profiling, since it allows for measurements that decouple the mutational fitness effects on folding and dynamics.

Comparing protein fitness following permutation with the contact energies at each position where protein termini were introduced revealed a correlation. Sites having high energetic frustration were more functionally tolerant to new termini created by permutation compared with low frustration sites. Permutations in the AMP binding domain, which has been selected for the highest contact energies, yielded variants that exhibit near WT levels of total activity in cells. Since the AMP binding domain is thought to control product release (12, 19), the rate limiting step of catalysis in AK, our findings suggest that the extra conformational flexibility created by permutation does not affect this mechanistic role. Additionally, our results suggest that the extra conformational flexibility arising from new termini in the AMP binding domain does not disrupt folding, substrate binding, or catalytic activity (Figure 6D). The variants with the lowest fitness across all three homologs arose from the creation of new protein termini at different locations within the core domain. Since these new termini were generated at locations that have selected for low contact energies, and previous biophysical studies have implicated the core domain as critical to overall protein thermostability (16), these findings suggest that increases in conformational entropy arising from permutation in this domain are deleterious to folding. Our discovery of permuted variants with fitness that exceeds the parental proteins suggests that these permuted AKs may differ in activity and/or folding from the parental AKs. In future studies, it will be interesting to characterize the biochemical and biophysical properties of these and other topological mutants discovered herein. Specifically, it will be interesting to investigate how the activity varies with temperature and whether the mechanism of product release is altered by permutation in the AMP binding and/or lid domain. Increased conformational flexibility in the AMP-binding domain caused by backbone cleavage may facilitate product release and improved enzyme turnover rates (12, 19). Additionally, it will be interesting to explore which of the topological mutants that are inactive retain structures that are similar to one of the conformational states sampled by the native proteins but are unable to navigate the full catalytic cycle because of changes in substrate binding or dynamics.

Our results indicate that CPP-seq is a powerful tool in protein engineering for scanning and identifying positions where increasing local entropy enhances function. This is counter to the prevailing approach of targeting increased native state stability through substitutions that improve packing (73) or decrease conformational entropy (74). Circular permutation scanning of AKs support the use of such strategies for their core domains, but not for positions proximal to the active site. A broad survey of high-resolution enzyme structures finds that active sites consistently exhibit higher energetic frustration than the rest of the protein (75), suggesting that increasing active site conformational entropy may be a general engineering strategy for enzymes beyond AK. Given that introducing chain termini in the middle of a protein sequence can cause structural and energetic perturbations other than increased local flexibility, *i.e.*, the larger size of N- and C-termini relative to the peptide bond, and the unfavorable desolvation energy of burying formally charged termini, it is quite possible that the fitness increases observed from near-active-site permutations may underestimate the potential for functional optimization at such positions through point mutations. In the future, it will be interesting to investigate whether CPP-seq can be used to identify positions where conformational entropy may be enhanced through substitutions to glycine or other small amino acids that reduce local packing.

## MATERIALS AND METHODS

### Data availability

Deep sequencing data are available from the NCBI Sequence Read Archive (SRA) under accession numbers SAMN13192615 (*Bg*-AK, unselected), SAMN13192616 (Bg-AK, selected), SAMN13192617 (Bs-AK, unselected), SAMN13192618 (Bs-AK, selected), SAMN13192619 (Gs-AK, unselected), SAMN13192620 (*Gs*-AK, selected). Python scripts used to analyze the data are available at github.com/SilbergLab. Analyzed data for figures is available upon request.

### Vector Construction

The genes encoding *Bacillus globisporus*, *Bacillus subtilis*, and *Geobacillus stearothermophilus* AK were PCR amplified from pNIC28-BgAK, pNIC28-BsAK, and pNIC28-GsAK to create amplicons flanked by NotI sites, the addition of a spacer base before the start codon to maintain in-frame translation and removal of the stop codon, cloned into pET26b vectors using Golden Gate assembly to yield pET26b-BgAK, pET26b-BsAK, and pET26b-GsAK. All vectors were sequence verified.

### Library construction

Each AK vector (~4 µg each) was digested with NotI overnight at 37 °C, then agarose gel electrophoresis was used to separate the AK genes from other reaction products, and each AK gene was purified using a Zymoclean^TM^ Gel DNA Recovery Kit (Zymo Research) and eluted with DNA-grade water. Purified AK genes (~400 ng) were circularized using T4 DNA ligase in a 20 µL reaction incubated at 16 °C for 16 hours. Following ligation, circularized AK genes were further purified using a DNA Clean & Concentrator Kit (Zymo Research) and eluted with 12 µL of DNA-grade water. The yield of each circularized AK was assessed using a NanoDrop spectrophotometer (220-350 ng). To create each library, the circular genes were mixed with BglII linearized pMT-P1 (50-150 ng), an artificial transposon with all of the attributes of a protein expression vector (Addgene #120863), and 1 unit of MuA transposase (Thermo Fisher Scientific) in a 20 µL reaction and incubated at 37°C for 16 hours. Following incubation at 75°C for 10 minutes, total DNA from each reaction was purified using a DNA Clean & Concentrator kit and transformed (~200-400 ng) into library grade MegaX DH10B Ultracompetent cells (Thermo Fisher Scientific) using electroporation. Following electroporation, cells were allowed to recover in recovery media (Thermo Fisher Scientific) for 45 minutes at 37°C while shaking before plating 200 µL of recovery culture onto five LB-agar plates (150 × 15 mm) containing 25 µg/mL of kanamycin. After incubation at 37°C for 24 hours, CFU were quantified visually. Estimates of unselected library sampling based on colony counts on plates indicated sampling exceeded the number of possible variants (1320) by >20 fold. To harvest the final libraries, 3 mL of LB was added to each plate, a sterile spreader was used to homogenize colonies, the cell slurries from each library were pooled, and a QIAprep Spin Miniprep Kit (Qiagen) was used to isolate each unselected library. The library construction reaction yielded three types of DNA, circularized permuteposon (pMT2), permuteposon-AK gene hybrids in the parallel orientation, and permuteposon-AK gene hybrids in the antiparallel orientation (Figure S8). The band corresponding to the permuteposon-AK hybrids was cut out from the agarose gel, and this DNA was purified using a Zymoclean^TM^ Gel DNA Recovery Kit and DNA-grade water elution. Restriction digest analysis of these purified size-selected libraries reveal that they were homogeneous for permuteposon-AK gene hybrids (Figure S9).

### Library selections

To quantify the fraction of functional AKs in each library, the purified unselected libraries (300 ng) were transformed into *E. coli* CV2 using electroporation, cells were spread on LB-agar plates containing 25 µg/mL kanamycin, and five plates containing each library were incubated at 30 °C and 42 °C for 48 hours. Visual inspection of these plates was used to estimate total CFU, and the fraction functional was calculated as the number of CFU at 42°C divided by the number at 30°C. The number of unique selected CFU in each library was also estimated by dividing the observed CFU by two to account for the doubling of cells that occurs during the growth recovery following transformation. Among the three libraries, this analysis revealed that we sampled 60,080 BgAK, 104,720 BsAK, and 48,200 GsAK variants. The colonies from each library were harvested and pooled, and a QIAprep Spin Miniprep Kit was used to isolate each selected library. The incubation times were determined by performing controls with *E. coli* CV2 transformed with a circularized transposon lacking an AK gene. When these cells were grown on LB-agar plates containing 25 µg/mL kanamycin at 42°C for 48 hours, no colonies were observed.

### Library sequencing

Each unselected and selected library (≥700 ng) was digested with restriction endonucleases that cut ~750 base pairs upstream of the permuted genes (ClaI) and ~150 base pairs downstream of the permuted gene (PciI). Agarose electrophoresis was used to separate the permuted AK genes from the vector backbone, and the bands encoding each ensemble of permuted AK genes (~1.6 kb) were excised and purified. All downstream processing and sequencing of the unselected and selected libraries was performed by the Baylor College of Medicine Genomic and RNA Profiling Core. A Nextera XT kit was used to fragment the DNA ensemble in each library and attach unique sequencing adapters to the ends. An Illumina MiSeq System was then used to collect single-end sequencing reads (150 base pairs) on all of the libraries in parallel.

### Sequence analysis

Sequencing data was analyzed using a previously described custom Python pipeline, which is available at github.com/SilbergLab/CPP-seq. In brief, this pipeline first identifies reads containing a permuteposon using nine base pair sequences at the end of each permuteposon. The pipeline then determines if each read is at the beginning or end of the permuteposon by analyzing for the presence of a larger 54 base pair sequence. Reads including the start codon are designated “start motif” reads, while those including the stop codon are designated the “stop motif” reads. Each of these reads is then further divided into sense and antisense reads. We next evaluate the first 11 base pairs of AK-derived sequence adjacent the start/stop motifs and determined how they related to all possible 11 base pair sequences within each of the AK genes. This comparison allows us to then designate AK genes as parallel (P) and in an orientation that can be transcribed and translated, or antiparallel (AP) and in an orientation that does not allow for expression.

### Statistical analysis

To analyze whether vectors were significantly enriched by the selection, we took a similar approach to the DESeq method (76). DEseq was developed to analyze if there are significant changes in gene expression with RNA-seq and ChIPseq datasets, using no change as the frame of reference and the null hypothesis. With our dataset, we used the ratio of selected to unselected AP variant counts as a frame of reference for the expected effect of selective pressure as the null hypothesis instead of the no change null hypothesis that DESeq assumes. We modeled the AP variant counts (i = 1 to 223 for each unique variant) using a negative binomial model where we defined X_i_ and Y_i_ as the unselected and selected counts, respectively. In our model, the AP counts in the unselected library are defined as m_i_ and the AP counts in the selected library are m_i_ multiplied by a scalar dilution factor (β), such that X_i_ = m_i_, Y_i_ = βm_i_, Var (X_i_) = γm_i_, and Var (Y_i_) = γβm_i_. There are two global parameters shared by all the loci, β and γ. The former reflects the average selection effect on all the AP variants, and the latter reflects the overdispersion compared with Poisson distribution. β was estimated using sample median, and γ was estimated by assuming a negative binomial distributions for X_i_ and Y_i_ and then maximizing the likelihood. Using the estimated negative binomial distribution, the P-value of each P variant pair (X_i_, Y_i_) was computed by conditioning on the sum of X_i_ and Y_i_ as outlined in equation eleven of DESeq (76). Finally, P-values were adjusted for multiple testing using the Benjamini–Hochberg procedure.

### Fitness calculations

To relate the relative activity of all three AK orthologs we converted the log_2_(fold change) into a measure of fitness relative to the native AK. This was done by normalizing each Fold Change by the Fold Change of the native AK present in the same library according to:

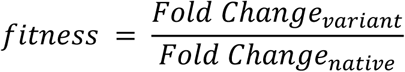

## ACKNOWLEDGEMENTS

We are grateful to Dr. George Phillips and Jose L. Olmos for their gift of the plasmids pNIC28-BgAK, pNIC28-BsAK, and pNIC28-GsAK. We are grateful for financial support from the National Science Foundation, including MCB grant number 1150138 to J.J.S. and Graduate Research Fellowships to A.M.J. and J.T.A. Additionally, this work was supported by Office of Naval Research grant N00014-17-1-2639 (to J.J.S.). AMJ was partially funded by a training fellowship from the Keck Center of the Gulf Coast Consortia, on the Houston Area Molecular Biophysics Program, National Institute of General Medical Sciences (NIGMS) T32GM008280. JTA was partially supported by a Lodieska Stockbridge Vaughn Fellowship. We thank Kevin R. MacKenzie, Peter Wolynes, George N. Phillips for helpful discussions.

## AUTHOR CONTRIBUTIONS

A.M.J., J.T.A., and J.J.S. designed the study. A.M.J. and J.T.A. performed experiments. J.T.A. analyzed the data. J.T.A., V.N., and J.J.S. wrote the manuscript

## SUPPLEMENTAL DATA

**Figure S1.**
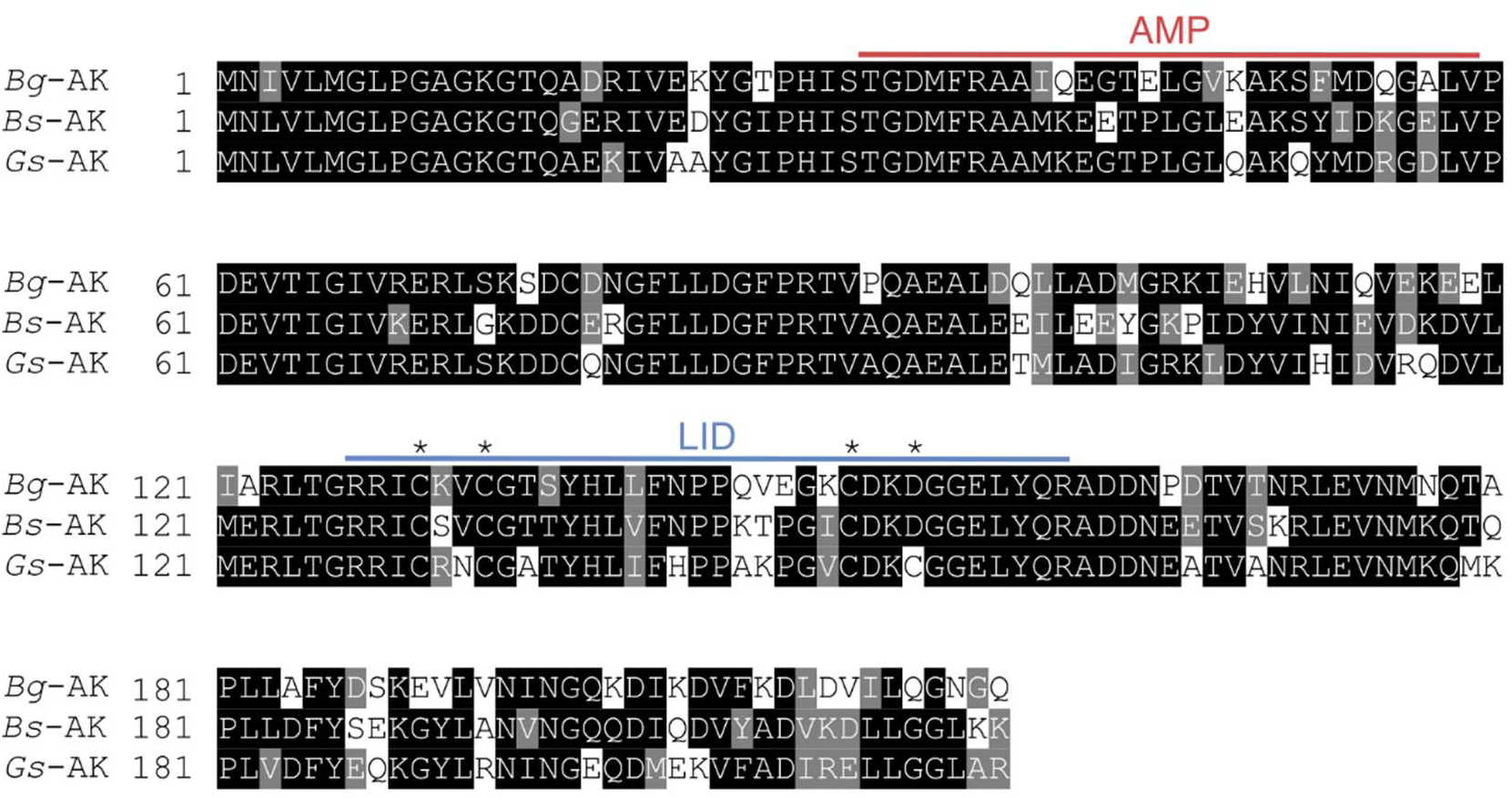
Multiple sequence alignment of adenylate kinases. A multiple sequence alignment generated by aligning *Bg*-AK, *Bs*-AK, and *Gs*-AK using MUSCLE (1). All three AKs are the same length (217 residues). Residue positions conserved across 50% of the sequences in the MSA are highlighted in black if identical and grey if similar amino acids. Residues from the mobile domains are shown in red (AMP binding), and blue (lid). The Cys-X_2_-Cys-X_16_-Cys-X_2_-Cys/Asp zinc-binding motif that is conserved in gram-positive bacterial AKs are marked with asterisk.

**Figure S2.**
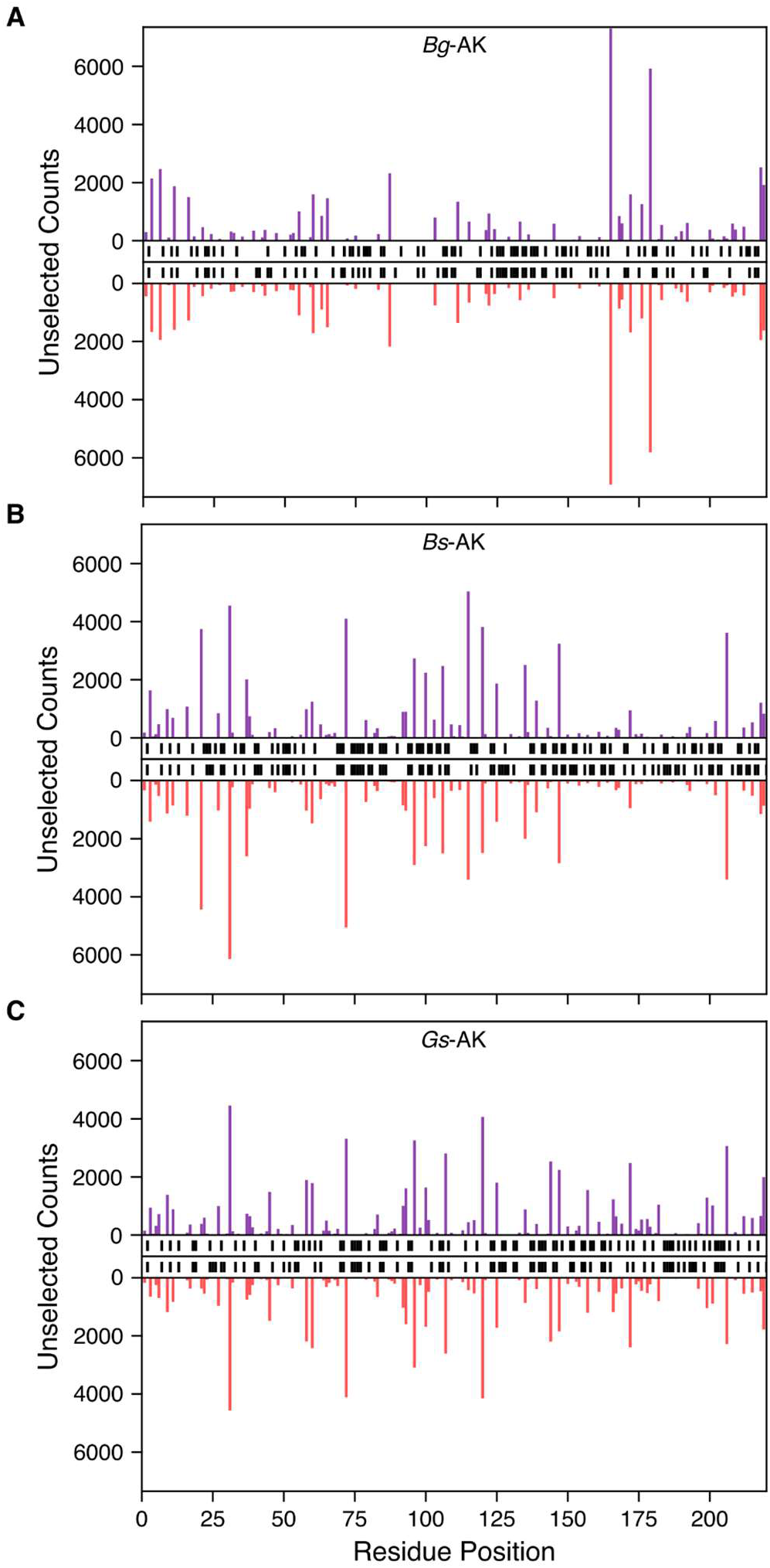
Sequence abundances as a function of the first AK codon in each ORF. A comparison of the AK codon at the beginning of each permuted gene and the number of P (purple) and AP (red) sequence reads observed in the unselected libraries for (*A*) *Bg*-AK, (*B*) *Bs*-AK, and (*C*) *Gs*-AK. In cases where a P or AP variant was absent, black bars are shown between the x-axes.

**Figure S3.**
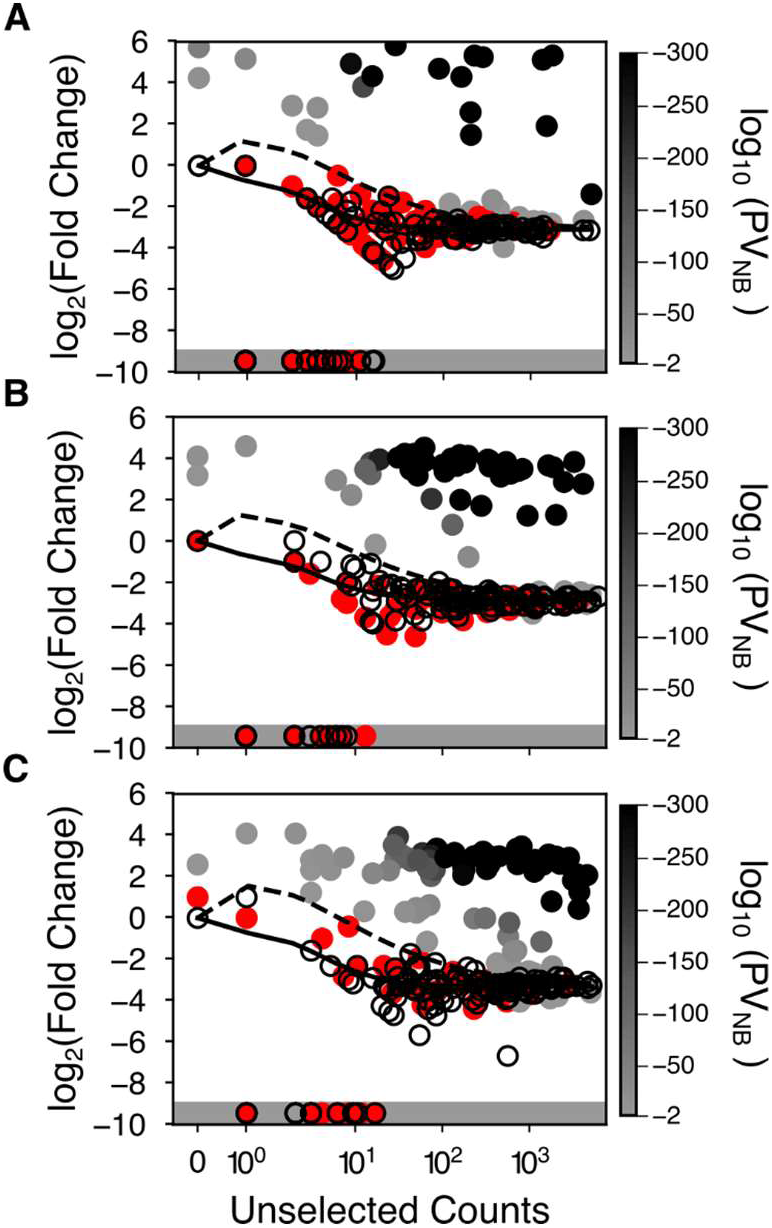
Fold enrichment as a function of the initial abundance and statistical model. The log_2_(fold change) in sequence abundances of the AP (open symbols) and P variants (closed symbols) for (*A*) *Bg*-AK, (*B*) *Bs*-AK, and (*C*) *Gs*-AK. The significance of P variant enrichment obtained using the negative binomial model (PV_NB_) is colored as a function of the *P*-value obtained with the variants presenting values >0.01 in red, variants having values ≤10^−300^ in black and those variants displaying intermediate values shaded as indicated in the bar. The black line represents the mean dilution for AP variants relative to their initial abundance in the unselected library, while the dashed line represents two standard deviations greater than the mean. Variants not observed in the selected library (infinitely diluted) are plotted in the shaded region.

**Figure S4.**
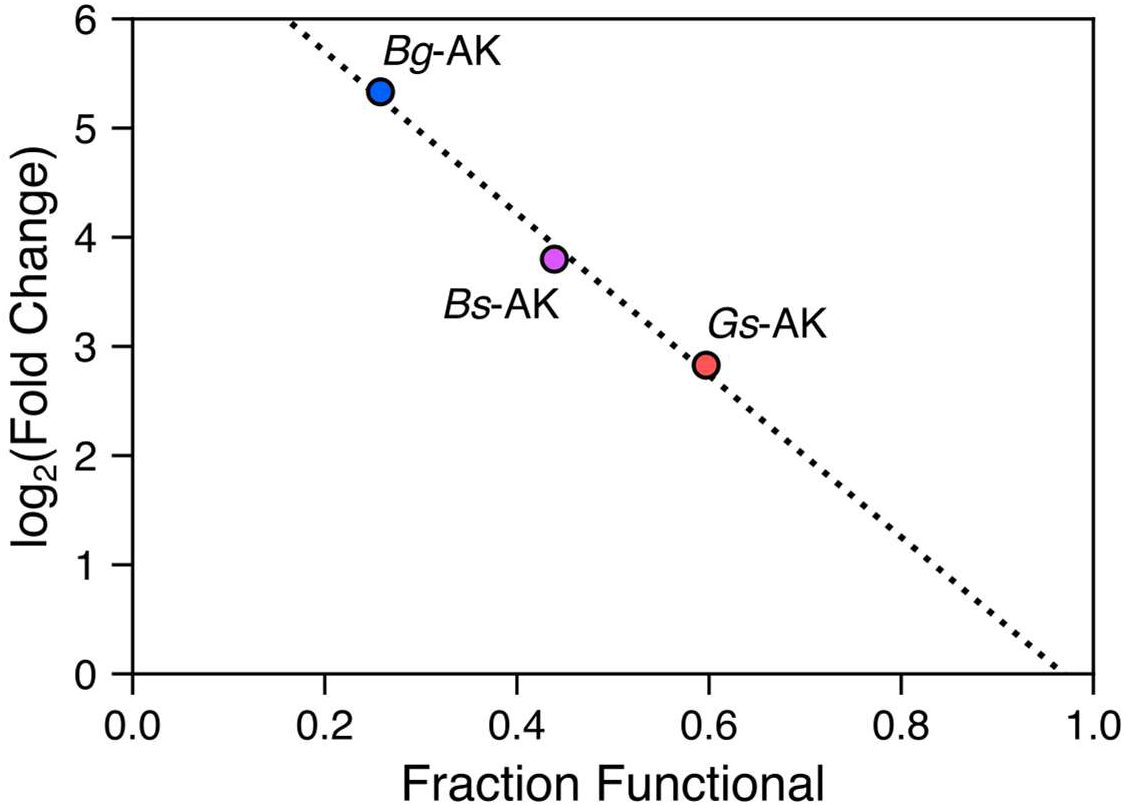
Relationship between fold change of native AK sequence reads and the fraction of functional variants. The fold change of the native AK in each library is compared with the fraction of P variants sampled in the libraries that were biologically active following selection. A linear fit (y = −7.41x - 7.18, R^2^ = 0.992) is shown as a dashed line.

**Figure S5.**
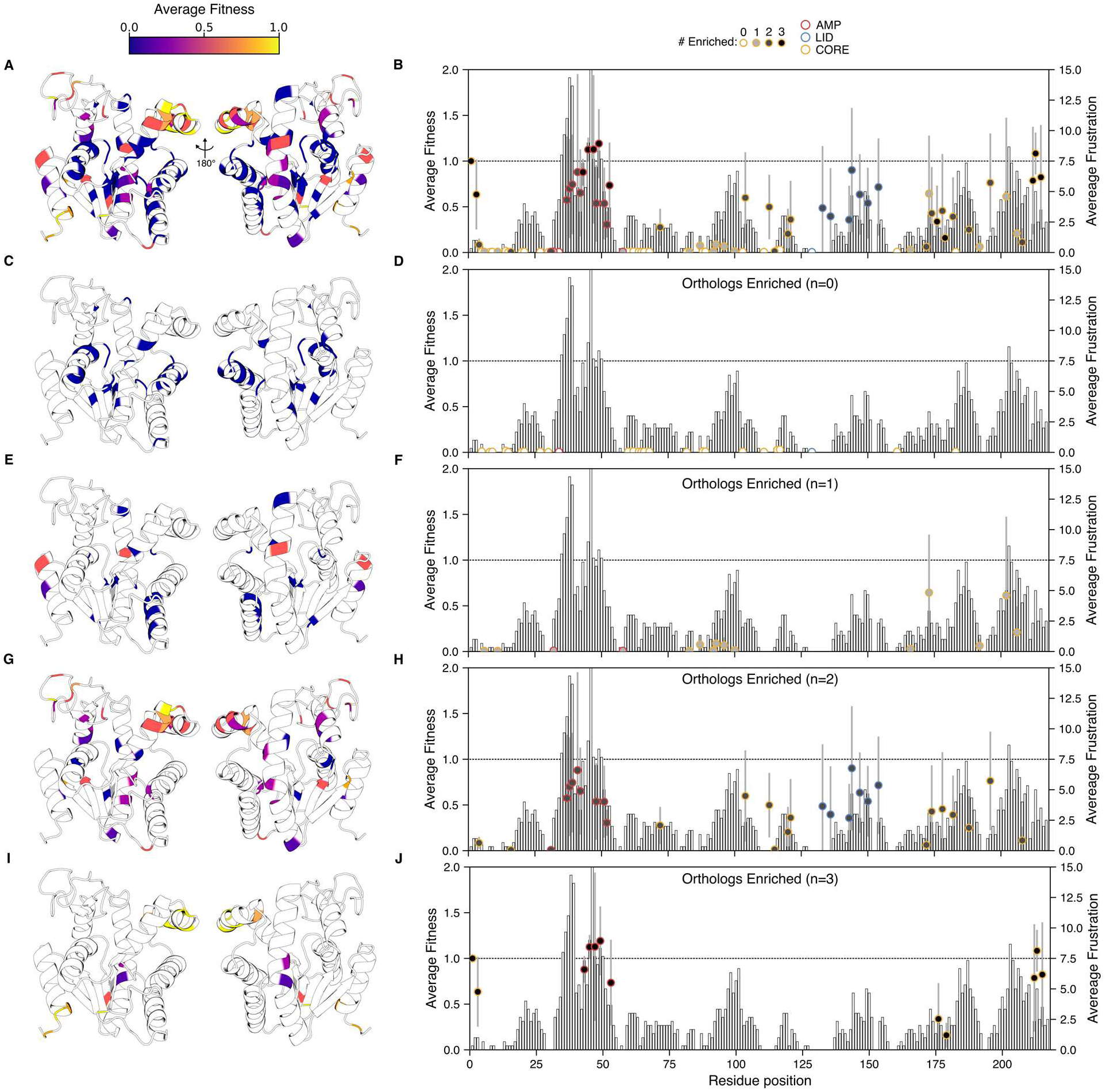
Comparison of family permutation profiles and energetic frustration. (*A*) The average fitness of each trio of topological mutant orthologs is compared to the tertiary structure (PDB: 1S3G). The color of the residue corresponds to the average fitness of a permutation at each site according to the color bar. Positions that were unobserved in ≥1 orthologs are shaded white. (*B*) The average fitness is compared to the position in the primary structure. The s.d. of each average is shown in gray. The shading of each circular data symbol represents the number of orthologs in which the variant was found to be significantly enriched. The bars represent the average density of highly frustrated contacts at each position. The dotted line represents the native AK fitness observed in each library. (*C*) and (*D*) show the same data for trios of AK orthologs that were uniformly inactive. (*E*) and (*F*) show the same data for trios of AK orthologs where only 1 out of 3 ortholog displayed cellular activity. (*G*) and (*H*) show the same data for trios of AK orthologs where 2 out of 3 orthologs presented cellular activity. (*I*) and (*J*) show the trios of AK orthologs where 3 out of 3 orthologs exhibited cellular activity.

**Figure S6.**
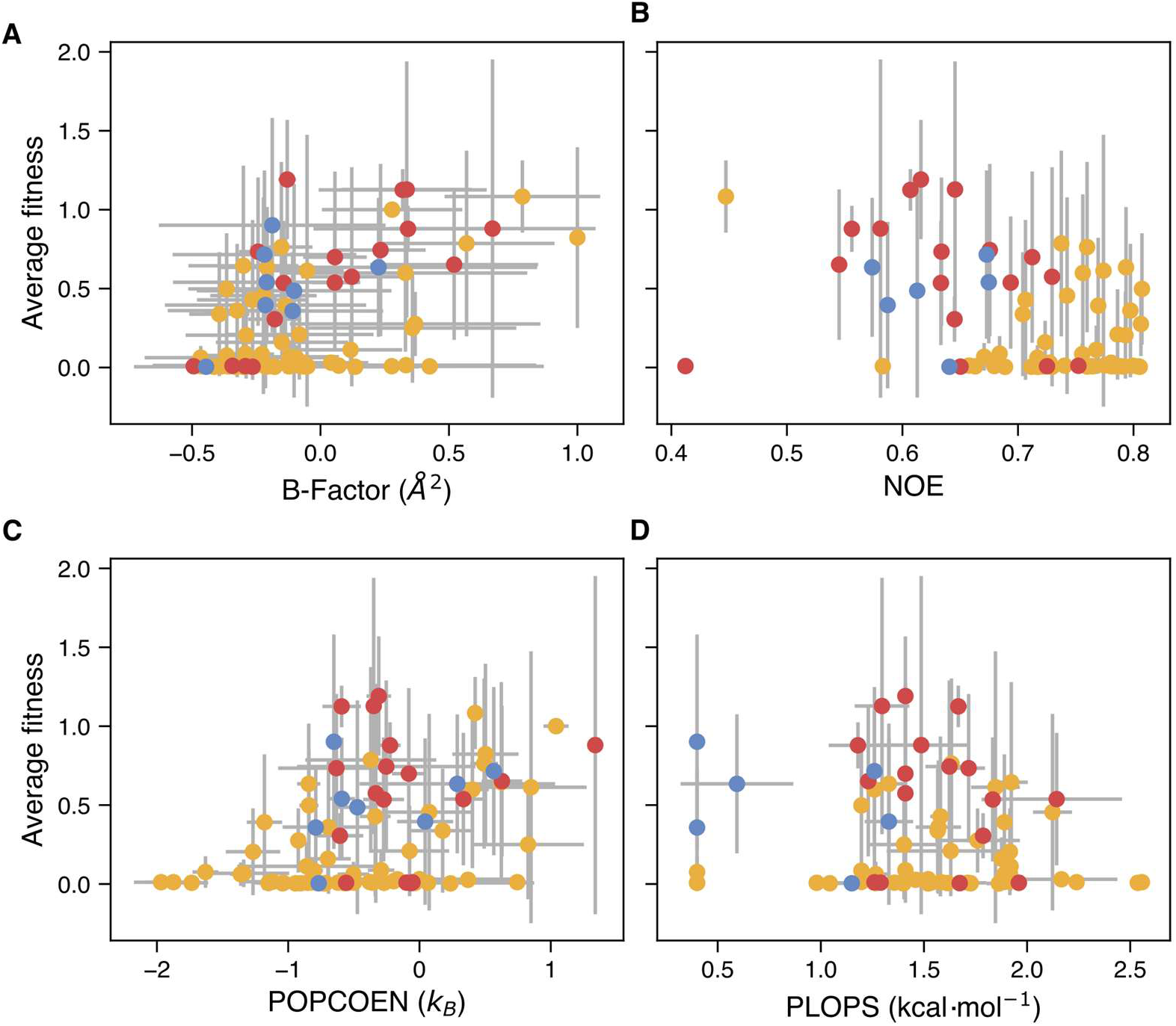
Comparison of protein dynamics metrics with the average fitness of permuted proteins. The average fitness (fold change relative to native AK) of each trio of orthologous permuted proteins is compared to (*A*) the average normalized B-factors from each crystal structure (R*_SR_*=0.439, *P*-value = 2.95 × 10^−5^), (B) the NOE data from *Ec*-AK (2) (R*_SR_*=-0.324, *P*-value = 6.00 × 10^−3^), (C) predictions of residue-level conformational entropy resulting from POPCOEN (3) (R*_SR_*=0.380, *P*-value = 3.71 × 10^−4^), and (D) prediction of residue-level conformational entropy loss upon folding resulting from PLOPS (4) (R*_SR_*=0.015, *P*-value = 8.96 × 10^−1^). B-factors were normalized using a previously described method (5)

**Figure S7.**
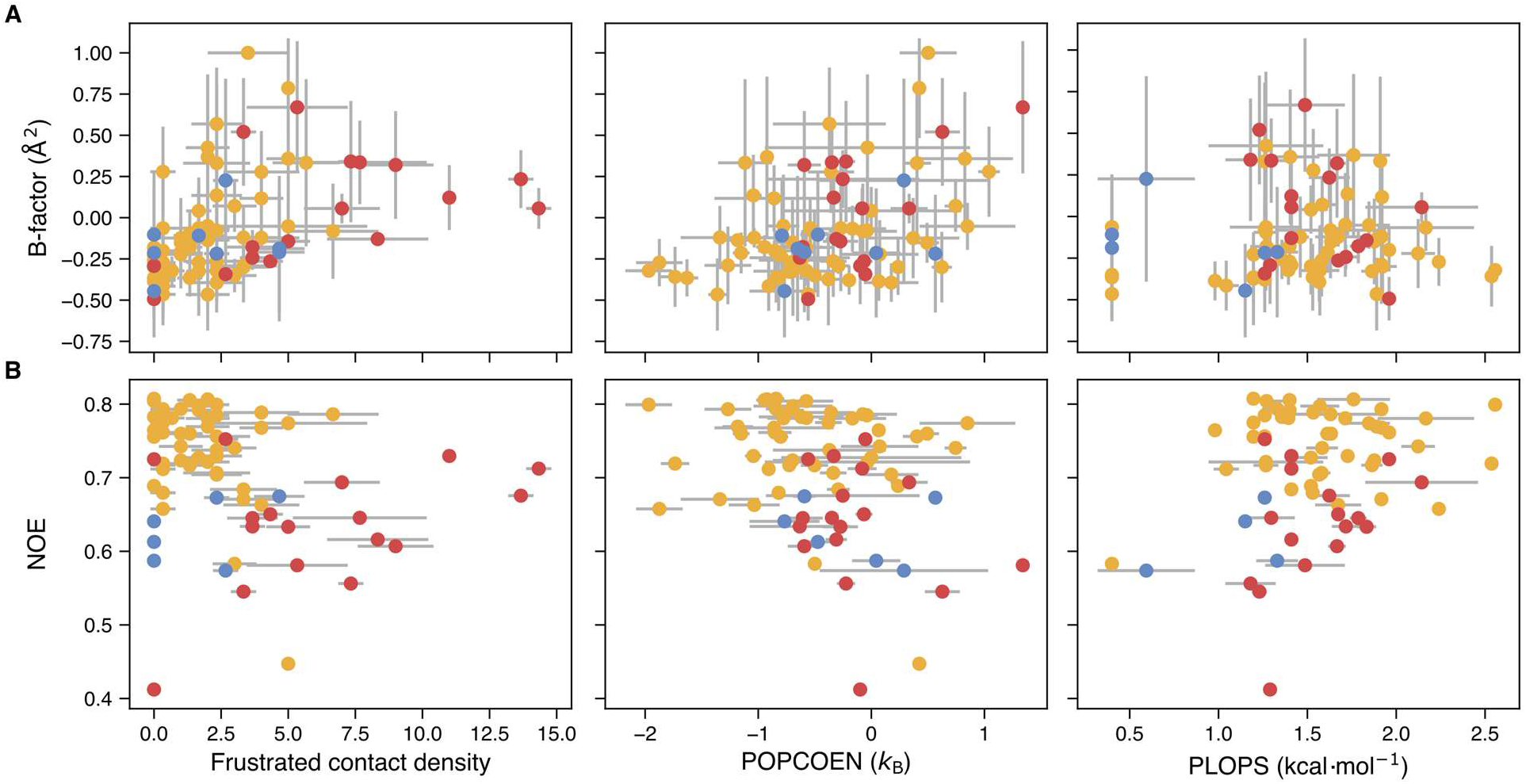
Comparison of experimental measures of protein dynamics with computational metrics. The pairwise comparison of (A) B-factors and (B) NOE data from *Ec*-AK is compared with predictions of the frustrated contact density calculated using the frustratometer, predictions of residue-level conformational entropy calculated using POPCOEN (3), and PLOPS predictions of residue-level conformational entropy loss upon folding (4). Correlations between the metrics are as follows: B-Factor and frustrated contact density (R*_SR_*=0.573, *P*-value = 1.22 × 10^−8^), B-factor and POPCOEN (R*_SR_*=0.348, *P*-value = 1.18 × 10^−3^), B-factor and PLOPS (R*_SR_*=0.073, *P*-value = 5.24 × 10^−1^), NOE and frustrated contact density (R*_SR_*=-0.315, *P*-value = 7.95 × 10^−3^), NOE and POPCOEN (R*_SR_*=-0.316, *P*-value = 7.76 × 10^−3^), and NOE and PLOPS (R*_SR_*=0.133, *P*-value = 2.87 × 10^−1^)

**Figure S8.**
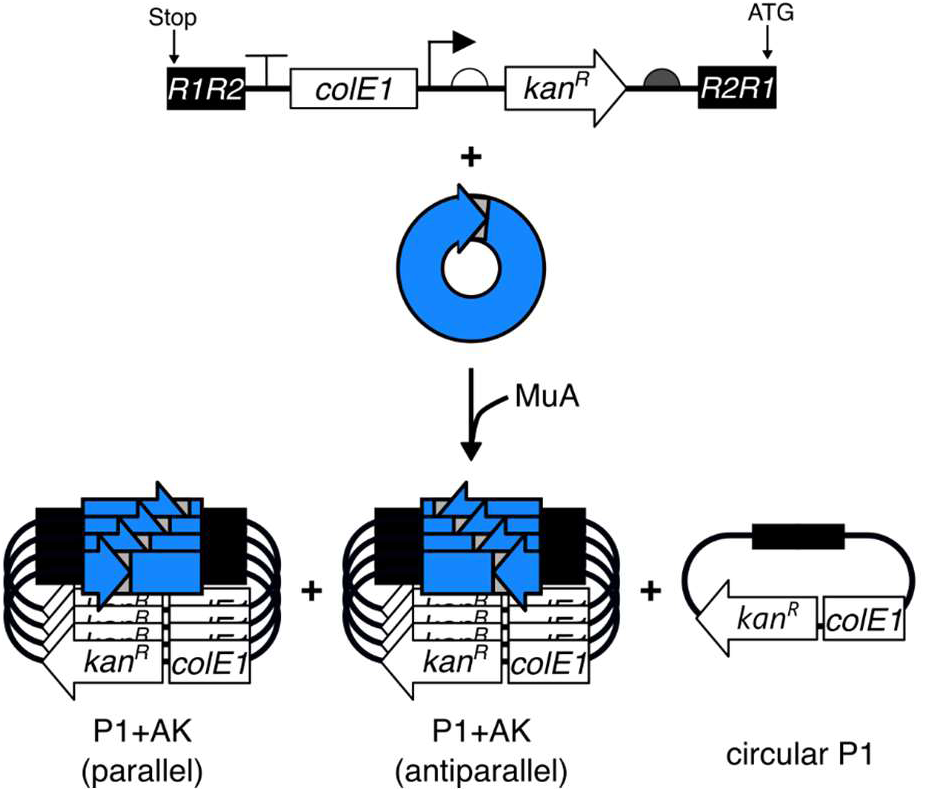
Analysis of permuted MuA libraries. (*A*) PERMUTE protocol uses MuA transposase to insert a permuteposon randomly into a circularized gene, which eliminates the need for using a target vector. The product of this reaction yields circularized permuteposon (P1) and the desired permuteposon + AK gene hybrids (P1 + AK) in both the parallel and antiparallel orientations.

**Figure S9.**
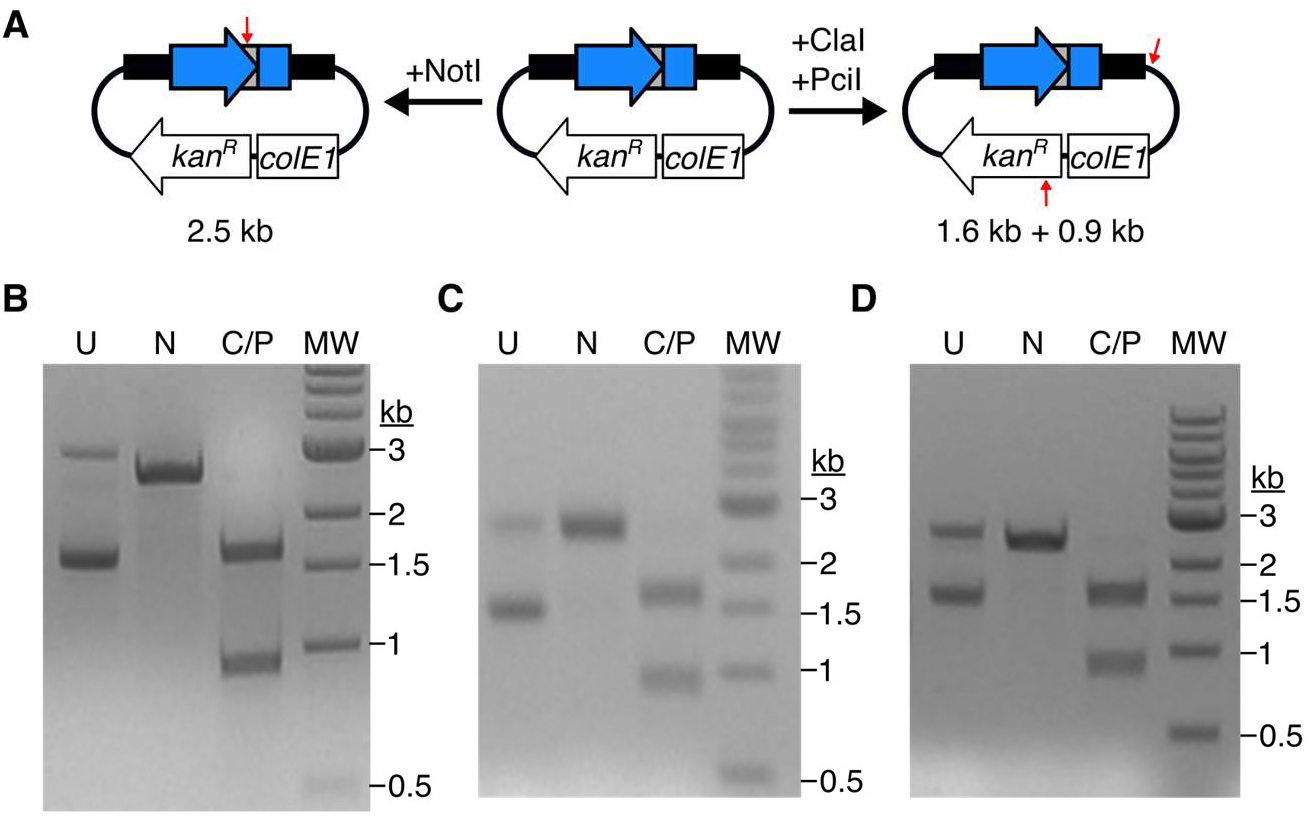
Size-selected permuted libraries are homogeneous for P1 + AK hybrids. (A) Following purification of the P1 + AK hybrids from the MuA libraries, the size-selected libraries were subjected to NotI (N) digests expected to yield a 2.5 kb fragment and ClaI/PciI (C/P) double digests expected to yield 1.6 and 0.9 kb fragments. Restriction site locations are marked by red arrows. Agarose gel electrophoresis of the digests reveals the correct bands in (*B*) *Bg*-AK, (*C*) *Bs*-AK, (*D*) *Gs*-AK libraries compared to the uncut MuA library control (U).

